# Endothelial neuropilin-1 and neuropilin-2 are essential for tumour angiogenesis

**DOI:** 10.1101/2022.05.17.492307

**Authors:** CJ Benwell, RT Johnson, JAGE Taylor, CA Price, SD Robinson

**Affiliations:** Gut Microbes and Health, Quadram Institute Bioscience, Norwich Research Park, Norwich, UK; School of Pharmacy, University of East Anglia, Norwich Research Park, Norwich, UK; School of Biological Sciences, University of East Anglia, Norwich Research Park, Norwich, UK

## Abstract

Neuropilin (NRP) expression is highly correlated with poor outcome in multiple cancer subtypes. As known co-receptors for vascular endothelial growth factor receptors (VEGFRs), core drivers of angiogenesis, past investigations have alluded to their functional roles in facilitating tumorigenesis by promoting invasive vessel growth. Despite this, it remains unclear as to whether NRP1 and NRP2 act in a synergistic manner to enhance pathological angiogenesis. Here we demonstrate, using NRP1^*ECKO*^, NRP2^*ECKO*^ and NRP1/NRP2^*ECKO*^ mouse models, that maximum inhibition of primary tumour development and angiogenesis is only achieved when both endothelial NRP1 and NRP2 are targeted simultaneously. Metastasis and secondary site angiogenesis were also significantly inhibited in NRP1/NRP2^*ECKO*^ animals. Mechanistic studies revealed that co-depleting NRP1 and NRP2 in mouse-microvascular endothelial cells (ECs) stimulates rapid shuttling of VEGFR-2 to Rab7^+^ endosomes for proteosomal degradation. Our results highlight the importance of targeting both NRP1 and NRP2 to modulate tumour angiogenesis.

## Introduction

Angiogenesis is a critical driver of tumour growth and metastatic dissemination. Without the expansion of a vascular network to supply oxygen and nutrients to the tumour, growth cannot proceed past a few millimetres [1]–[3]. Vascular endothelial growth factor (VEGF)- dependent stimulation of vascular endothelial growth factor receptor-2 (VEGFR-2) represents a major signalling pathway promoting angiogenesis, yet the clinical benefits of targeting the VEGF/VEGFR-2 axis remain modest. Only minimal increases in progression-free survival rates for various tumour types, including lung, breast, kidney, and colon cancers, have been reported following treatment [3]. Only when combined with chemotherapy have such therapies become recognised as an effective strategy against cancer growth, anti-angiogenics acting to selectively prune leaky, and immature tumour-associated vessels to facilitate more efficient delivery of chemotherapeutic agents [4]–[6]. The identification of novel combinations of angiogenic targets to enhance the therapeutic index of anti-VEGF/VEGFR-2 strategies remains paramount however, on account of cancers developing numerous adaptive mechanisms to escape tumour therapy [7].

Neuropilin 1 (NRP1) and 2 (NRP2) are type-1 transmembrane glycoprotein co-receptors for VEGFs and their respective VEGFRs [8], [9]. Notably, NRP1 is known to form complexes with VEGF-A_165_, a principle pro-angiogenic factor, and VEGFR-2 to promote angiogenic signalling, whilst NRP2 preferentially binds VEGF-C and VEGFR-3 to propagate lymphangiogenic signalling [3], [10], [11]. That said, NRP2 has also been shown to induce optimal thresholds of VEGFR-2 phosphorylation by promoting VEGFR-2/VEGF-A_165_ interactions, subsequently enhancing both survival and migratory signalling cascades [12]. Perhaps unsurprisingly, neuropilin (NRP) over-expression is often considered synonymous with an enhanced rate of tumour growth, invasiveness and angiogenesis in a number of different cancer types, including carcinoma [13], colorectal [14], melanoma [15], myeloid leukaemia [16], breast [17] and lung cancer [18]. This is facilitated, at least in part, by their ability to interact with integrin receptors, for example integrin α5β1, to enhance tumour cell spreading and extravasation [19], [20].

Owing to their ability to associate with a diverse range of receptors, in turn forming holoreceptors to propagate a plethora of downstream pro-angiogenic signalling cascades, NRPs are promising targets for anti-tumour therapies [3]. For example, the NRP1-specific small molecule inhibitors *EG00229* and *ATWLPPR* have been demonstrated to inhibit NRP1-VEGFR-2 signalling, and impair both tumour angiogenesis and tumour growth *in vivo* [21]– [23]. Tandem-virtual screening and cell-based screening have since been utilised by Borriello *et al*., to identify a series of non-peptide VEGF-NRP antagonists, notably NRPα-47 and NRPα-308, which display anti-angiogenic and anti-proliferative capabilities *in vitro*, in addition to anti-tumorigenic effects on breast cancer *in vivo* [24], [25]. More recently, the anti-tumour potential of NRPα-308 was employed against clear cell Renal Cell Carcinoma (ccRCC), a highly vascularised cancer arising from the overexpression of VEGF-A_165_. Compared to the tyrosine kinase inhibitor sunitinib, the current reference treatment for ccRCC, NRPα-308 was found to suppress ccRCC cell proliferation, migration and invasiveness to a greater extent. Gen depletion studies supported these findings, and alluded to the fact that both NRP1 and NRP2 should be completely inhibited to obtain maximal therapeutic effect [26].

Presently, there have been limited studies comparing the anti-angiogenic effects of depleting either NRP receptor individually versus when they are targeted together. To this end, we generated genetically modified mouse models that enabled us to perform temporal endothelial-specific deletions of either NRP gene individually, or in combination. Utilising these models, we demonstrate that in multiple models of cancer, co-targeting the endothelial expression of both NRP1 and NRP2 severely inhibits primary and secondary tumour growth and angiogenesis, to a much greater extent than when either NRP receptor is targeted alone. The depletion of both NRP1 and NRP2 severely impairs fibronectin containing extra domain-A (EDA-FN) secretion *in vivo* and *in vitro*, which likely impedes pathological vessel stability and growth. We also demonstrate that NRP depletion stimulates the rapid degradation of VEGFR-2, metering surface receptor availability for VEGF-A_165_-induced pro-angiogenic responses.

## Results

### Dual targeting of endothelial expressed NRPs inhibits primary tumour growth and angiogenesis by impairing vessel stability

As co-receptors for VEGF family receptors, endothelial neuropilins (NRPs) are becoming increasingly recognised as candidate targets for supressing pathologies typified by uncontrolled vascular expansion, such as cancer and retinopathy. Investigations have, however, persisted in elucidating their function separately from one another, rather than in conjunction. For example, by crossing NRP2-floxed (NRP2^flfl^) mice [27] with mice expressing a tamoxifen-inducible Pdgfb-iCreER^T2^ promoter [28], we previously showed that endothelial NRP2 (NRP2^*EC*^) promotes pathological angiogenesis to support the progression of primary tumours in a lung-carcinoma model. Acute, endothelial-specific depletion of NRP2 impaired tumour development and vascularisation approximately 2-fold, revealing a novel, effective therapeutic strategy against cancerous growth [29]. Indeed, Kaplan-Meier plots indicate a significantly reduced overall patient survival following diagnosis with lung carcinoma when either NRP1 or NRP2 mRNA expression is elevated [30] (**Suppl. Figure 1A-B**).

To ascertain whether endothelial NRP1 and NRP2 contribute in a non-redundant, synergistic manner during angiogenesis-dependent tumour development, we crossed NRP2^flfl^.Pdgfb-iCreER^T2^ (NRP2^flfl^.EC^KO^) mice with NRP1^flfl^.Pdgfb-iCreER^T2^ (NRP1^flfl^.EC^KO^) mice to generate NRP1^flfl^;NRP2^flfl^.Pdgfb-iCreER^T2^ (NRP1^flfl^NRP2^flfl^.EC^KO^) animals and compared the effects of an acute endothelial-specific depletion of NRP1, NRP2 or NRP1;NRP2 during subcutaneous allograft tumour growth using CMT19T lung carcinoma cells. Tamoxifen administrations were performed thrice weekly starting 4 days prior to CMT19T cell implantation, continuing until day 18 (D18), at which point primary tumours were harvested (**Figure 1A**). NRP1^flfl^.EC^KO^ and NRP2^flfl^.EC^KO^ animals developed significantly smaller tumours (∼50%) compared to their respective Pdgfb-iCreER^T2^-negative (Pdgfb-iCreER^T2-^) controls, as observed previously [29]. In comparison, tumours harvested from NRP1^flfl^NRP2^flfl^.EC^KO^ mice grew significantly smaller than either NRP1^flfl^.EC^KO^ or NRP2^flfl^.EC^KO^ tumours, and only to ∼20% the size of those harvested from control mice (**Figure 1B-E**). No changes to gross animal weight were observed (**Suppl. Figure 1C**). Immunofluorescence imaging of endomucin^+^ blood vessels verified that our tamoxifen regimen effectively silenced target expression. Notably, NRP1^flfl^NRP2^flfl^.EC^KO^ tumours also exhibited significantly less vasculature than either NRP1^flfl^.EC^KO^ or NRP2^flfl^.EC^KO^ tumours (**Figure 1F-G**), suggesting that the dual targeting of both NRP1 and NRP2 elicits a compounded anti-angiogenic response to inhibit tumour development and growth.

**Figure 1:**
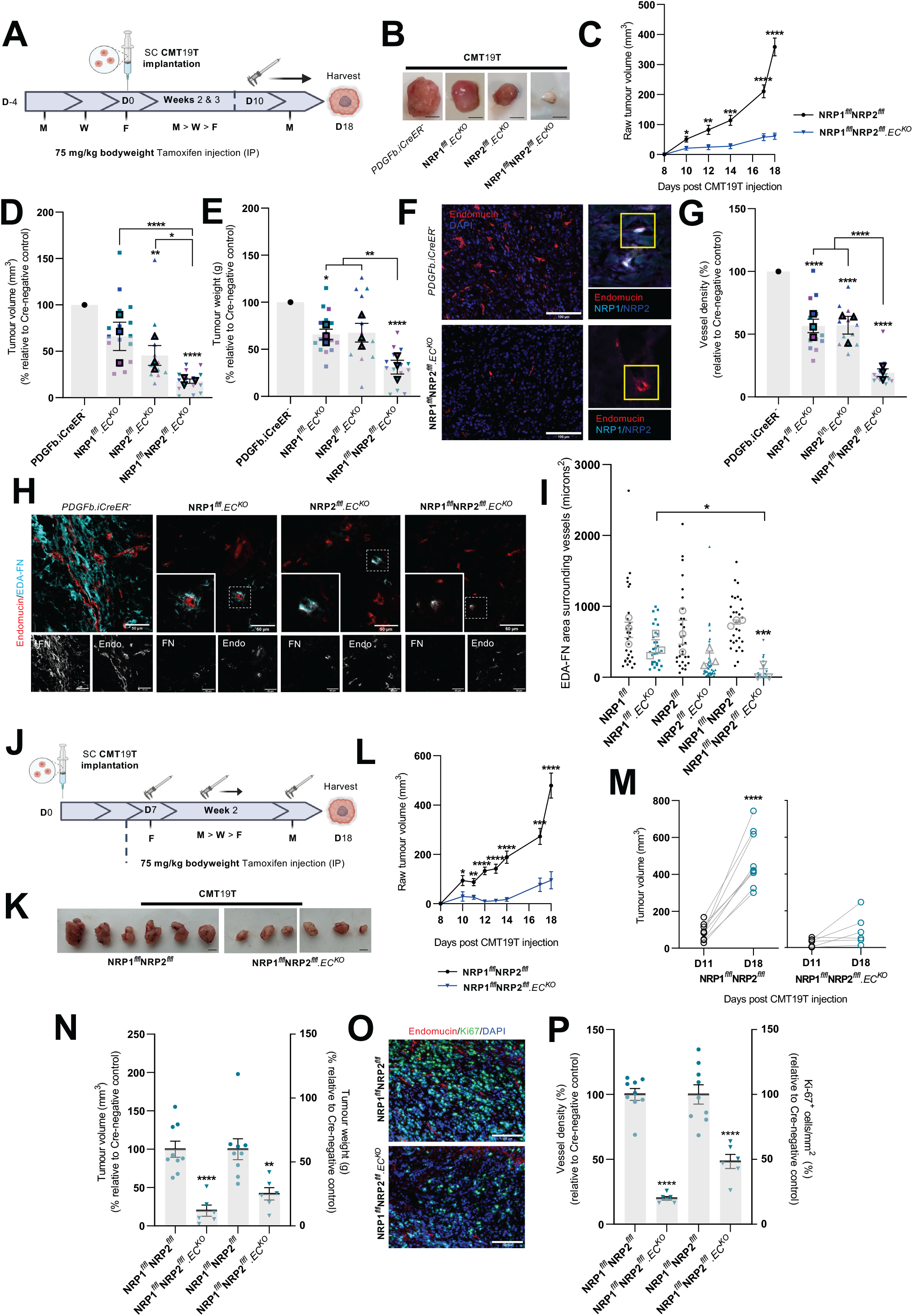
Dual targeting of endothelial expressed NRPs effectively inhibits primary tumour growth. Inducible, endothelial specific deletion of NRPs, either individually, or in combination was achieved by crossing mice expressing the PDGFb.iCreER promoter of Cre-recombinase to those floxed for NRP1, NRP2 or NRP1/NRP2. **A**) Experimental schematic: tamoxifen-induced activation of Cre-recombinase and thus deletion of targets was employed via the following regime. Cre-positive and Cre-negative littermate control mice received intraperitoneal (IP) injections of tamoxifen (75 mg/kg bodyweight, 2mg/ml stock) thrice weekly (Monday, Wednesday, Friday) for the duration of the experiment from D-4 to D17 to induce Cre-recombinase activity. CMT19T lung carcinoma cells (1×10^6^) were implanted subcutaneously (SC) into the flank of mice at D0 and allowed to grow until D18. **B**) Representative images of CMT19T tumours harvested on D18 removed from Cre-negative and positive mice. Scale bar shows 5 mm. **C**) Raw tumour volume growth kinetics from 10 days post CMT19T injection to harvest. Tumour volume calculated using the formula: length x width^2^ x 0.52. Error bars show mean ± SEM; N=3 (n≥12). **D**) Quantification of relative tumour volumes measured on D18. Data presented as a percentage of the average tumour volume (mm^3^) observed in their Cre-negative littermate controls. Error bars show mean ± SEM; N=3 (n≥12). **E**) Quantification of tumour weight (g) measured on D18. Data presented as percentages of the average tumour weight (g) observed in respective littermate controls. Error bars show mean ± SEM; N=3 (n≥12). **F**) (*Left panels*) Representative tumour sections from Cre-negative and Cre-positive tumours showing endomucin^+^ blood vessels. Scale bar = 100 µm. (*Right panels*) Confirmation of endothelial-specific target depletion in tumour sections from Cre-negative and Cre-positive tumours. **G**) Quantification of % blood vessel density per mm^2^. Mean quantification performed on 3x ROIs per tumour section, from 1-3 sections per tumour. Data presented as a percentage of the average % vessel density observed in their Cre-negative littermate controls. Error bars show mean ± SEM; N=3 (n≥12). **H**) Representative tumour sections from Cre-negative and Cre-positive tumours showing EDA-FN coverage around endomucin^+^ blood vessels. Scale bar = 100 µm. **I**) Quantification of mean EDA-FN area (µm^2^) surrounding vessels, performed on ≥10 vessels/tumour. Error bars show mean ± SEM; n≥3. **J**) Delayed experimental schematic: tamoxifen-induced activation of Cre-recombinase and thus deletion of targets was employed via the following regime. Cre-positive and Cre-negative littermate control mice received intraperitoneal (IP) injections of tamoxifen (75 mg/kg bodyweight, 2mg/ml stock) thrice weekly (Monday, Wednesday, Friday) from D7 to induce Cre-recombinase activity. CMT19T lung carcinoma cells (1×10^6^) were implanted subcutaneously (SC) into the flank of mice at D0 and allowed to grow until D18. **K**) Representative images of CMT19T tumours harvested on D18 removed from Cre-negative and positive mice. Scale bar shows 5 mm. **L**) Raw tumour volume growth kinetics from 10 days post CMT19T injections to harvest. Error bars show mean ± SEM; n≥6. **M**) Raw tumour volume growth change measured between D11 and D18, n≥6. **N**) Quantification of tumour volume (mm^3^) (*Left axis*) and weight (g) (*Right axis*) measured on D18. Data presented as percentages of the average tumour volume and weight observed in respective littermate controls. Error bars show mean ± SEM; n≥6. **O**) Representative tumour sections from Cre-negative and Cre-positive tumours showing endomucin^+^ blood vessels and Ki-67^+^ proliferating cells. Scale bar = 100 µm. **P**) Quantification of % blood vessel density per mm^2^ (*Left axis*) and % number of Ki-67^+^ proliferating cells per mm^2^ (*Right axis*) from CMT19T tumours. Mean quantification performed on 3x ROIs per tumour section, from 1-3 sections per tumour. Data presented as a percentage of the average % vessel density/% number of Ki-67^+^ proliferating cells observed in their Cre-negative littermate controls. Error bars show mean ± SEM; n≥6. Asterixis indicate significance. **Panels A-G**: Permission granted by FASEB journal [29] to reuse some data under the terms of the Creative Commons CC BY license.

The extracellular matrix (ECM) component fibronectin containing extra domain-A (EDA-FN) is a known marker of tumour vasculature, and is essential for the development of a metastatic microenvironment [20], [31]–[33]. As both NRP1 and NRP2 have been reported to regulate FN fibrillogenesis in ECs in the past [29], [34], we considered whether the deposition of EDA-FN around tumour vessels would be perturbed in our knockout models. Compared to respective Cre-negative control tumours, only those depleted for both NRP1 and NRP2 saw a significant reduction in EDA-FN coverage around tumour vasculature (**Figure 1H-I**), suggesting that both endothelial NRPs facilitate tumour angiogenesis by promoting vessel stability. Deoxycholate studies employing siRNA-transfected wild-type (WT) mouse-lung ECs validated these findings, combined depletion of both NRP1 and NRP2 resulted in a significant reduction in EDA-FN expression from both unstimulated and VEGF-A_165_-stimulated insoluble fractions (**Suppl. Figure 1D-E**).

To determine if co-targeting endothelial NRP1 and NRP2 impedes tumour growth in already established tumours, we next performed intervention CMT19T allograft studies in our NRP1^flfl^NRP2^flfl^.EC^KO^ animals whereby we delayed tamoxifen administrations until 7 days after cell implantation (**Figure 1J**). By doing so, we aimed to provide a more clinically relevant study design, where treatment is initiated once a cancer has become vascularised. Following this regimen, we observed severe impediments to tumour growth and angiogenesis following the combined loss of endothelial NRP1 and NRP2. Tumours in NRP1^flfl^NRP2^flfl^.EC^KO^ mice were significantly smaller than controls or NRP1^flfl^.EC^KO^ or NRP2^flfl^.EC^KO^ animals (**Figure 1K-N**). Again, no changes in mean animal weight were observed between groups **(Suppl. Figure 1F**). In addition to a significant reduction in tumour vascularity, NRP1^flfl^NRP2^flfl^.EC^KO^ tumours also exhibited significantly fewer Ki67^+^ proliferating cells compared to control tumours (**Figure 1O-P**).

Finally, to exclude tumour size as a statistical confounder, and therefore assess whether the loss of endothelial NRP1 and NRP2 directly influences pathological angiogenesis, tamoxifen administration was suspended further until day 12 (**Suppl. Figure 1G**). Tumour growth was tracked from day 7, and tumours were harvested from all animals on day 16. Whilst no significant reduction in tumour volume was detected between NRP1^flfl^NRP2^flfl^.EC^KO^ mice and Cre-negative control mice, (**Suppl. Figure 1H-I**), immuno-labelling of endomucin-positive vessels revealed a ∼50% reduction in tumour vascularity (**Suppl. Figure 1J-K**). These results strongly suggest that co-depletion of endothelial NRP1 and NRP2 influences tumour angiogenesis regardless of tumour size, and in already highly vascularised tumours.

### Primary tumour development and angiogenesis is susceptible to the effects of co-targeting endothelial NRP1 and NRP2 in multiple cancer models

As the endothelial co-depletion of NRP1 and NRP2 was found to effectively impair primary lung carcinoma growth and angiogenesis, we proceeded to assess the efficacy of their co-depletion in other paradigms of cancer. To investigate whether the loss of NRP expression influences melanoma development, B16-F10 melanoma cells were subcutaneously implanted and allowed to grow for a period of 18 days, following our intervention-based tamoxifen regime as previously described in **Figure 1I** (**Figure 2A**). From D11, NRP1^flfl^NRP2^flfl^.EC^KO^ tumours grew significantly smaller than their control counterparts, and when excised were found to have developed to only ∼10% the size of control tumours. Indeed, a small number of tumours were found to have regressed entirely (**Figure 2B->D**). No changes in mean animal weight were observed between control and NRP1^flfl^NRP2^flfl^.EC^KO^ mice (**Figure 2E**).

**Figure 2:**
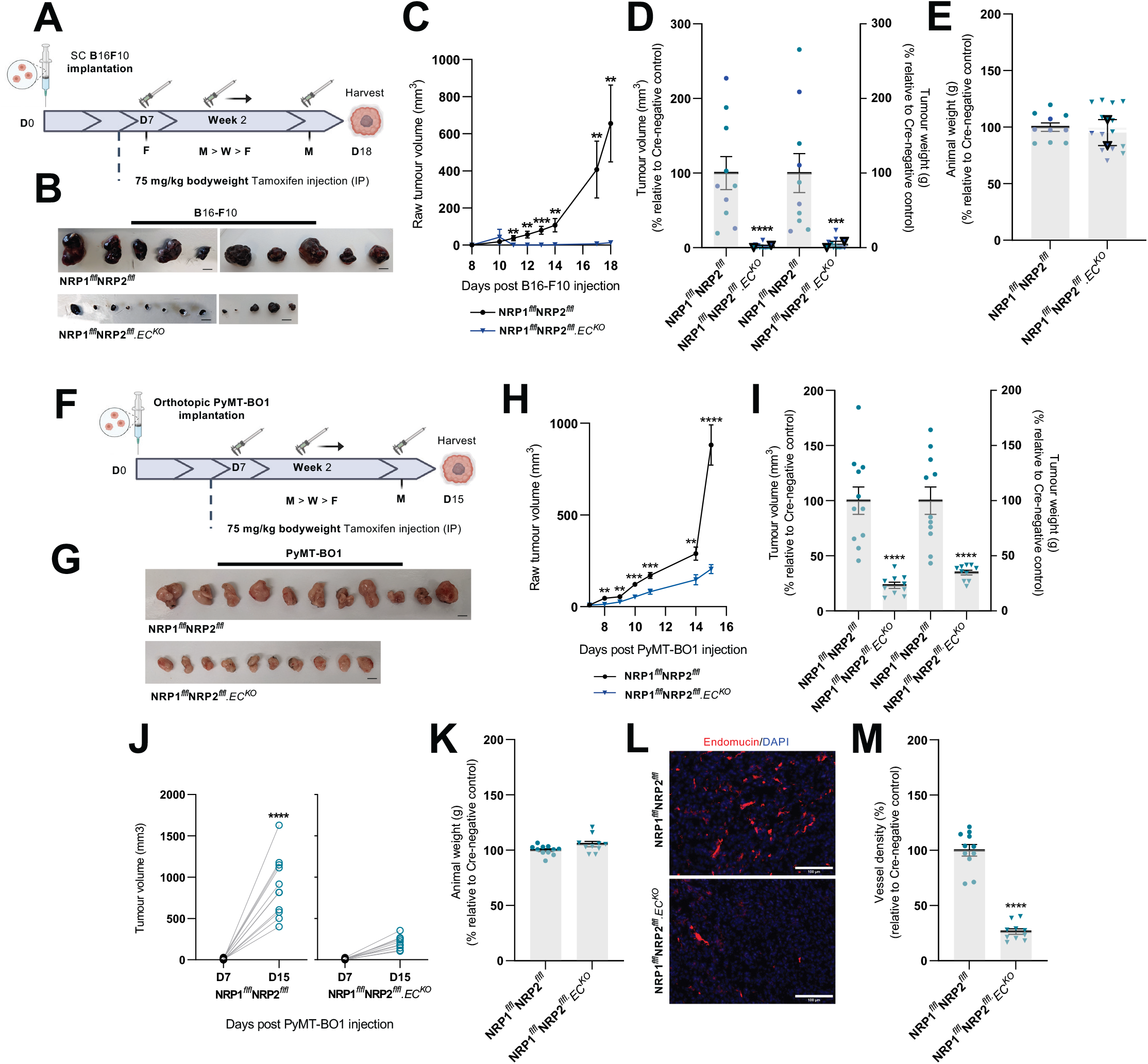
Primary tumour development and angiogenesis are susceptible to the effects of co-targeting endothelial NRP1 and NRP2 in multiple cancer models. Inducible, endothelial specific deletion of NRPs, either individually, or in combination was achieved by crossing mice expressing the PDGFb.iCreER promoter of Cre-recombinase to those floxed for NRP1/NRP2. **A**) Experimental schematic: tamoxifen-induced activation of Cre-recombinase and thus deletion of targets was employed via the following regime. Cre-positive and Cre-negative littermate control mice received intraperitoneal (IP) injections of tamoxifen (75 mg/kg bodyweight, 2mg/ml stock) thrice weekly (Monday, Wednesday, Friday) from D7 to induce Cre-recombinase activity. B16F10 melanoma cells (4×10^5^) were implanted subcutaneously (SC) into the flank of mice at D0 and allowed to grow until D18. **B**) B16F10 tumours harvested on D18 removed from Cre-negative and positive mice. Scale bar shows 5 mm. **C**) Raw tumour volume growth kinetics from 10 days post B16F10 injection to harvest. Tumour volume calculated using the formula: length x width^2^ x 0.52. Error bars show mean ± SEM; N=2 (n≥10). **D**) Quantification of tumour volume (mm^3^) (*Left axis*) and weight (g) (*Right axis*) measured on D18. Data presented as percentages of the average tumour volume and weight observed in respective littermate controls. Error bars show mean ± SEM; N=2 (n≥10). **E**) Quantification of mean animal weight measured at point of harvest. Data presented as a percentage of the average animal weight observed in respective littermate controls. Error bars show mean ± SEM, N=2 (n≥10). **F**) Experimental schematic: Cre-positive and Cre-negative littermate control mice received intraperitoneal (IP) injections of tamoxifen (75 mg/kg bodyweight, 2mg/ml stock) thrice weekly (Monday, Wednesday, Friday) from D7 to induce Cre-recombinase activity. PyMT-BO1 breast cancer cells (1×10^5^) were implanted orthotopically into the flank of mice at D0 and allowed to grow until D15. **G**) PyMT-BO1 tumours harvested on D15 removed from Cre-negative and positive mice. Scale bar shows 5 mm. **H**) Raw tumour volume growth kinetics from 10 days post PyMT-BO1 injection to harvest. Tumour volume calculated using the formula: length x width^2^ x 0.52. Error bars show mean ± SEM; n≥10. **I**) Quantification of tumour volume (mm^3^) (*Left axis*) and weight (g) (*Right axis*) measured on D15. Data presented as percentages of the average tumour volume and weight observed in respective littermate controls. Error bars show mean ± SEM; n≥10. **J**) Raw tumour volume growth change measured between D7 and D15, n≥10. **K**) Quantification of mean animal weight measured at point of harvest. Data presented as a percentage of the average animal weight observed in respective littermate controls. Error bars show mean ± SEM, n≥10. **L**) Representative tumour sections from Cre-negative and Cre-positive tumours showing endomucin^+^ blood vessels. Scale bar = 100 µm. **M**) Quantification of % blood vessel density per mm^2^ from PyMT-BO1 tumours. Mean quantification performed on 3x ROIs per tumour section, from 1-3 sections per tumour. Data presented as a percentage of the average % vessel density observed in their Cre-negative littermate controls. Error bars show mean ± SEM; n≥10. Asterixis indicate significance.

In a similar manner, we assessed the impact of co-depleting endothelial NRP1 and NRP2 on a luminal B model of breast cancer. PyMT-BO1 cancer cells [35] were orthotopically implanted into the fourth inguinal mammary gland of NRP1^flfl^NRP2^flfl^ and NRP1^flfl^NRP2^flfl^.EC^KO^ mice, and allowed to grow over a period of 15 days. Again, tamoxifen administration was delayed until day 7 to allow for palpable tumours to develop prior to the onset of target depletion (**Figure 2F**). Compared to NRP1^flfl^NRP2^flfl^ control tumours, those grown in NRP1^flfl^NRP2^flfl^.EC^KO^ animals developed ∼65% smaller by day 15 (**Figure 2G->J**), alongside no significant alterations in mean animal weight (**Figure 2K**). Unlike the B16-F10 tumours, which failed to grow more than ∼2 mm in size in our NRP1^flfl^NRP2^flfl^.EC^KO^ mice, we were able to process our PyMT-BO1 tumours for immunofluorescence imaging analysis. NRP1^flfl^NRP2^flfl^.EC^KO^ PyMT-BO1 tumours were ∼70% less vascularised than respective Cre-negative control tumours (**Figure 2L->M**), corroborating our CMT19T studies, and confirming that the expression of NRP1 and NRP2 is essential for tumour angiogenesis in multiple cancer models.

### Endothelial NRP1 and NRP2 co-depletion reduces the metastatic potential of circulating melanoma cells

Not only are murine B16-F10 cells a well-established, aggressive tumour model for preclinical investigations into melanoma progression, but they are also known to preferentially metastasise to the lungs of C57/BL6 mice [36]. To investigate whether dual-targeting of endothelial NRPs is effective at supressing hematogenous metastasis, we measured pulmonary seeding 14 days post intravenous injection of luciferase^+^-tagged B16-F10 cells (**Figure 3A**).

**Figure 3:**
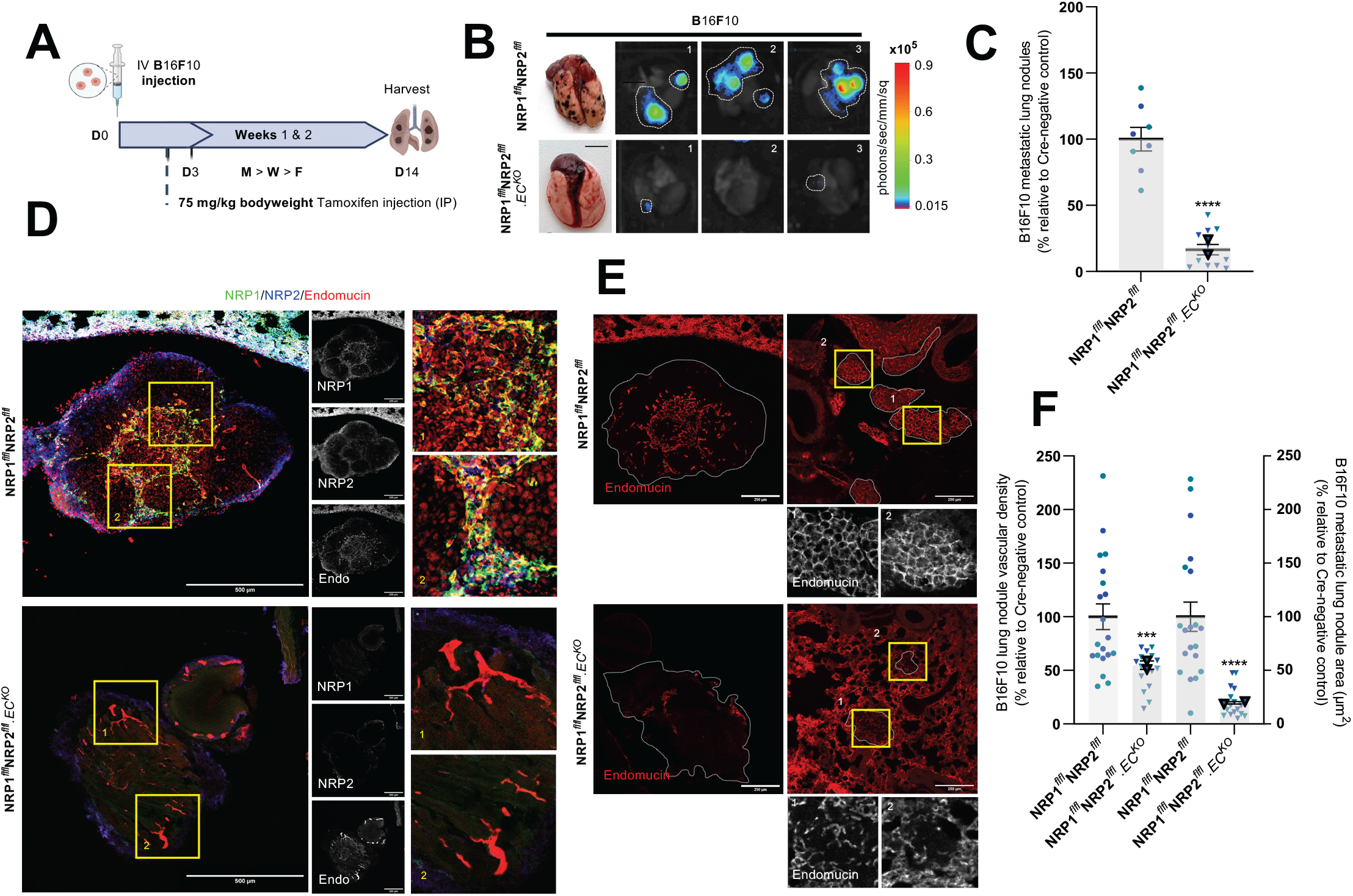
Endothelial NRP1 and NRP2 co-depletion reduces the metastatic potential of circulating melanoma cells. Inducible, endothelial specific deletion of NRPs, either individually, or in combination was achieved by crossing mice expressing the PDGFb.iCreER promoter of Cre-recombinase to those floxed for NRP1/NRP2. **A**) Experimental metastasis schematic: tamoxifen-induced activation of Cre-recombinase and thus deletion of targets was employed via the following regime. Cre-positive and Cre-negative littermate control mice received intraperitoneal (IP) injections of tamoxifen (75 mg/kg bodyweight, 2mg/ml stock) thrice weekly (Monday, Wednesday, Friday) from D3 to induce Cre-recombinase activity. B16F10 luciferase^+^ melanoma cells (1×10^6^) were intravenously (IV) injected into the tail vein of mice at D0 and allowed to disseminate until D14. **B**) (*Left panels*) Representative images of lungs harvested on D14 from Cre-negative and positive mice showing metastatic lung nodules. (Right panels) Corresponding representative bioluminescence (photons/sec/mm^2^) imaging of 3 lungs detected using Bruker imager. **C**) Quantification of number of metastatic nodules per lung at D14. Data presented as percentages of the average number of nodules observed in respective littermate controls. Error bars show mean ± SEM; N=2 (n≥8). **D**) Representative images of B16F10 metastatic nodules from Cre-negative and Cre-positive lungs showing NRP1, NRP2 and endomucin expression, and target knockdown. Scale bar = 250 µm. Boxed images show highlighted magnified regions. **E**) Representative images of B16F10 metastatic nodules from Cre-negative and Cre-positive lungs showing endomucin^+^ blood vessels. Scale bar = 250 µm. **F**) Quantification of nodule vascular density (*Left axis*) and nodule area (µm^2^) (*Right axis*). Data presented as percentages of the average vascular density and nodule area observed in respective littermate controls. Error bars show mean ± SEM; N=2 (n≥18). Asterixis indicate significance.

NRP1^flfl^NRP2^flfl^.EC^KO^ mice were found to develop significantly fewer metastatic lung nodules than control mice, subsequently confirmed by bioluminescence imaging (**Figure 3B->C**). Immunofluorescence staining of lung metastases revealed a robust expression of both NRP1 and NRP2 colocalising to endomucin^+^ vasculature in control nodules, but not in lung nodules of NRP1^flfl^NRP2^flfl^.EC^KO^ mice (**Figure 3D**). Furthermore, lung metastases of NRP1^flfl^NRP2^flfl^.EC^KO^ mice were observed to be significantly smaller and less vascularised than their control counterparts (**Figure 3E->F**). These results clearly demonstrate that the dual-targeting of endothelial NRP1 and NRP2 can be implemented not only as a means to retard primary tumorigenesis, but also to significantly reduce secondary site angiogenesis and growth.

### Endothelial NRPs regulate VEGFR-2 turnover to sustain pro-angiogenic signalling responses

Sustained hyperactivation of VEGFR-2 is largely considered one of the most critical aspects of pathological angiogenesis during tumour growth. Both NRP1 and NRP2 are also known co-receptors of VEGFRs and their respective VEGF signalling moieties [8], [9]. We therefore examined whether VEGFR-2 signalling is perturbed in tumour vasculature depleted for NRP1 and NRP2 by measuring VEGFR-2 and phosphorylated-VEGFR-2^Y1175^ localisation to endomucin^+^ vessels. Whilst NRP1^flfl^.EC^KO^ and NRP2^flfl^.EC^KO^ CMT19T tumours saw reductions in VEGFR-2 localisation of approximately 30% and 10% respectively, we observed a compounded reduction of over 50% in NRP1^flfl^NRP2^flfl^.EC^KO^ tumours (**Figure 4A->B**). Likewise, simultaneous depletion of both endothelial NRP1 and NRP2 resulted in an equivalent loss of phosphorylated-VEGFR-2^Y1175^ expression from tumour vessels (**Figure 4C, Suppl. Figure 2A**).

**Figure 4:**
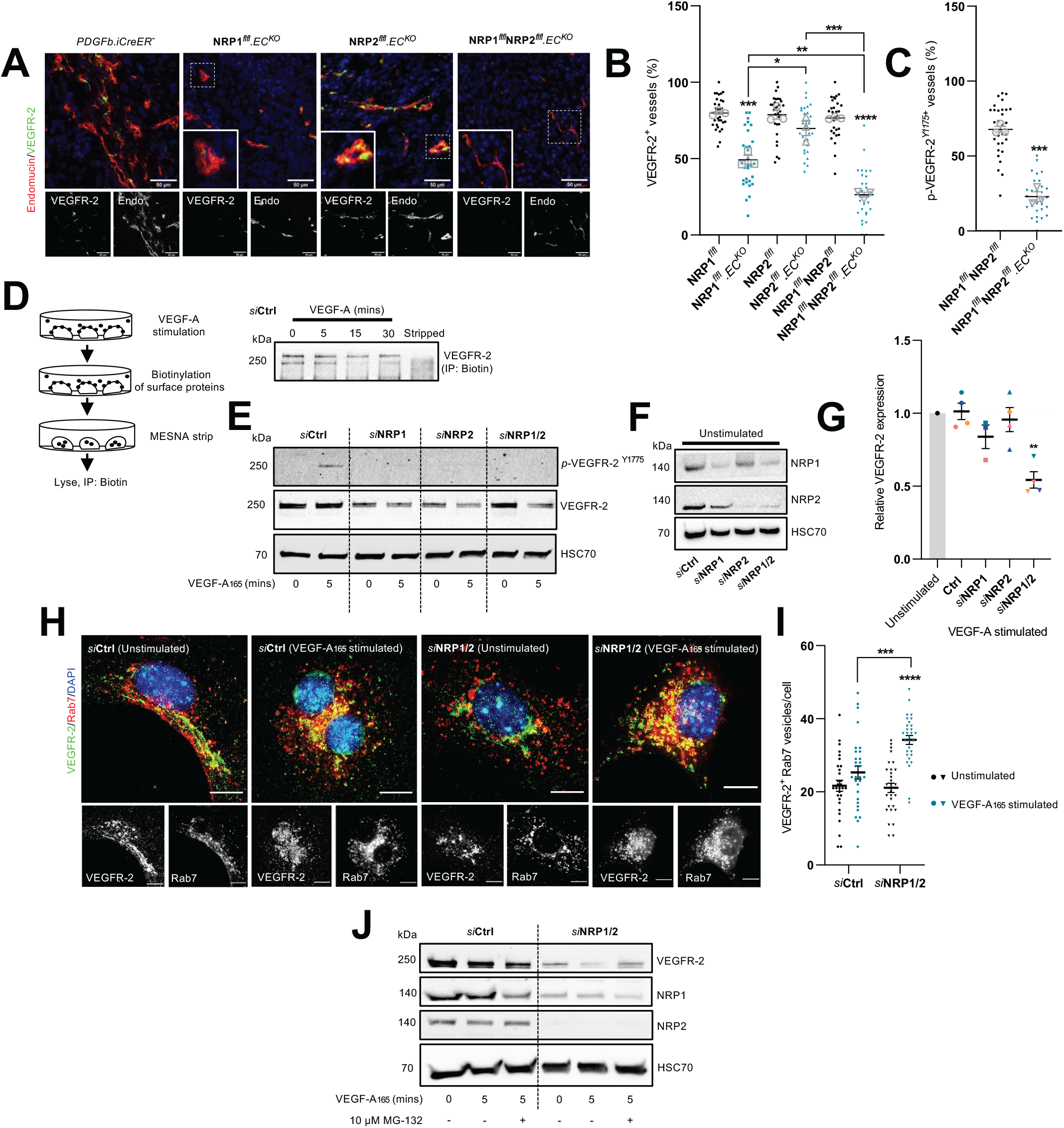
Endothelial NRPs regulate VEGFR-2 turnover to sustain pro-angiogenic signalling responses. **A**) Representative tumour sections from Cre-negative and Cre-positive CMT19T tumours showing colocalisation between VEGFR-2 and endomucin^+^ blood vessels. Scale bar = 100 µm. **B**) Quantification of % VEGFR-2^+^ vessels per mm^2^, performed on 10x ROIs per tumour. Error bars show mean ± SEM; n≥3. **C**) Quantification of % *p*-VEGFR-2^Y1175+^ vessels per mm^2^, performed on 10x ROIs per tumour. Error bars show mean ± SEM; n≥3. **D**) (Top panel) Schematic showing method for surface protein labelling and stripping. (Bottom panel) Ctrl siRNA-treated ECs were seeded onto 10 µg/ml FN for 48 hours before being incubated in serum-free OptiMEM for 3 hours. ECs were then stimulated with 30 ng/ml VEGF-A for the indicated timepoints before being labelled with 0.3 mg/ml biotin. Unreacted biotin was quenched with 100 mM glycine. ECs were then either lysed or incubated with 100 mM MESNA to strip off biotin labelled proteins. Unreacted MESNA was quenched with 100 mM iodoacetamide before lysis. EC lysates were immunoprecipitated with Protein G Dynabeads^TM^ coupled to anti-biotin primary antibody. Immunoprecipitated biotin-labelled proteins were separated by SDS-PAGE and subjected to Western blot analysis. Membranes were incubated in anti-VEGFR-2 primary antibody. **E**) siRNA-treated ECs were seeded onto 10 µg/ml FN for 48 hours, then incubated in serum-free OptiMEM for 3 hours. ECs were subject to 5 minutes of stimulation with 30 ng/ml VEGF-A before being washed twice on ice with PBS and lysed. Lysates were quantified using the DC protein assay, separated by SDS-PAGE and subjected to Western blot analysis. Membranes were incubated in anti-VEGFR-2, anti-*p*-VEGFR-2^Y1175^ and anti-HSC70 primary antibodies. **F**) Confirmation of target depletion by siRNA transfection. **G**) Quantification of total VEGFR-2 expression following VEGF-A stimulation relative to respective unstimulated lysates. Quantification shows mean densitometric analysis obtained using ImageJ^TM^. Error bars show mean ± SEM; N≥3. **H**) siRNA-treated ECs were seeded onto acid-washed, oven-sterilised coverslips pre-coated with 10 µg/ml FN for 3 hours. ECs were incubated in serum-free OptiMEM for 3 hours, before being subject to 5 minutes of stimulation with 30 ng/ml VEGF-A. Coverslips were fixed in 4% PFA, blocked and permeabilised. ECs were incubated with anti-VEGFR-2 and anti-Rab7 primary antibodies overnight at 4°C before incubation with appropriate Alexa fluor secondary antibodies at RT for 1 hour. Coverslips were mounted with flouromount G with DAPI^TM^. Panels show representative images of unstimulated and VEGF-A stimulated siRNA-treated ECs. Error bars show 10 µm. **I**) Quantification of VEGFR-2^+^ Rab7 vesicles/cell. Error bars show mean ± SEM; n=30. Asterixis indicate significance. **J**) siRNA-treated ECs were seeded onto 10 µg/ml FN for 48 hours, then incubated in serum-free OptiMEM ± 10 µM MG-132 for 3 hours. ECs were then stimulated, lysed and prepped for Western blot analysis in the same manner as **E**).

To further elucidate how endothelial NRPs co-operate to influence VEGFR-2 activity, we examined VEGFR-2 dynamics *in vitro*. First, we established VEGFR-2 surface expression levels in Ctrl siRNA-treated ECs remained intact up to 5 minutes after stimulation with VEGF-A_165_ (**Figure 4D**) by biotin labelling. Total cell lysates from Ctrl, NRP1, NRP2 and NRP1/2 siRNA-treated ECs were subsequently analysed by Western blotting to assess changes in VEGFR-2 expression following an acute 5 minute period of VEGF-A_165_ stimulation. This revealed that total VEGFR-2 expression was significantly diminished in stimulated ECs depleted for both NRP1 and NRP2 compared to unstimulated knockdown ECs (**Figure 4E-G**).

Following internalisation, VEGFR-2 is shuttled from Rab5^+^ early endosomes to either Rab4/Rab11^+^ recycling endosomes or rapidly degraded via Rab7^+^ late endosomes. We therefore proceeded to determine any changes in the fraction of VEGFR-2 localising to Rab7^+^ punctae following 5 minutes of VEGF-A_165_ stimulation. *si*NRP1/2 depleted ECs displayed a significantly greater proportion of VEGFR-2 present in Rab7^+^ vesicles compared to *si*Ctrl ECs (**Figure 4H-I**), suggesting that NRP1 and NRP2 promote VEGFR-2-induced pro-angiogenic responses by moderating receptor turnover. To test this hypothesis, we treated *si*Ctrl and *si*NRP1/2 ECs with 10 µM MG-132, a well characterised proteosome inhibitor [37]–[39]. MG-132 treatment effectively rescued total VEGFR-2 expression in VEGF-A_165_-stimulated *si*NRP1/2 depleted ECs (**Figure 4J**), confirming that NRP co-depletion stimulates the rapid translocation of VEGFR-2 from Rab7^+^ late endosomes to the proteosome for degradation.

## Discussion

Pathological angiogenesis is a core driver of aggressive tumorigenesis, yet the clinical benefits of targeting principle regulators of pro-angiogenic cascades have thus far shown limited efficacy [3]. We demonstrate that the endothelial-specific co-targeting of both NRP receptors, NRP1 and NRP2, provides effective inhibition against tumour growth and secondary site metastasis in multiple cancer models, likely by potentiating the rapid delivery of VEGFR-2 to late-endosomes for degradation. Importantly, we highlight the importance of targeting the expression of both NRPs simultaneously for maximum therapeutic effect.

Previous investigations have indeed demonstrated that targeting the expression of either NRP1 or NRP2 individually confers some anti-tumorigenic response. For example, inhibiting NRP1 binding to VEGF-A_165_ enhances the anti-tumour efficacy of VEGF-A_165_ blocking antibodies such as bevacizumab to modulate tumour cell proliferation and angiogenesis [40], [41]. The NRP1 inhibitor *EG00229* has also been demonstrated to exert significant tumour-suppressive effects in gliomas and squamous cell carcinomas [42]–[44]. Equally, treatment with the NRP2-specific monoclonal antibody *N2E4* inhibited pancreatic ductal adenocarcinoma (PDAC) cell tumour growth and metastasis by blocking interactions with β1 integrin to inhibit FAK/Erk/HIF-1α/VEGF-A_165_ signalling [45]. In addition, blocking NRP2 binding to VEGF-C was shown to reduce tumoral lymphangiogenesis and metastasis of breast adenocarcinoma and glioblastoma cells [46]. Naturally, it has since been elucidated that co-targeting the functions of both NRP1 and NRP2 may provide enhanced anti-tumorigenic responses [11].

Consistent with the above investigations, we confirm that an endothelial-specific deletion of either NRP gene significantly impairs tumour development and tumour angiogenesis. Critically however, dual loss of both NRP1 and NRP2 was found to reduce primary tumour growth and primary tumour angiogenesis by a greater extent than when either molecule was targeted individually. Furthermore, co-targeting NRP1 and NRP2 expression effectively inhibited secondary site metastasis compared to control animals. Given both NRP1 and NRP2 are known to modulate primary and secondary tumour microenvironments by interacting with integrins to remodel the tumoral ECM [20], [45], [47], of which FN is known as a major component [48], it follows that the impaired tumour growth exhibited following NRP co-depletion likely arises as a result of perturbations in EDA-FN fibril assembly and deposition. Indeed, EDA-FN has been demonstrated to facilitate tumour growth and invasiveness by promoting matrix stiffness, sustaining tumour-induced angiogenesis and lymphangiogenesis via VEGF-A_165_ [49] and VEGF-C respectively [50]. For example, Su *et al*., revealed that EDA-FN secretion promoted VEGFR-2 recruitment to β1 integrin sites, upregulating VEGFR-2 phosphorylation and pathological angiogenesis during hepatic fibrosis in a CD63-dependent manner [49].

As they are canonical co-receptors for VEGFR-2 in endothelial cells, we hypothesised that NRP co-depletion would provide effective inhibition of VEGFR-2-induced responses. NRP1/NRP2 knockout tumour vasculature was found to express significantly less VEGFR-2 than either NRP1 or NRP2 knockout tumours, in addition to reduced phosphorylated VEGFR-2 expression. Mechanistic studies utilising siRNA transfected WT mouse-lung ECs subsequently revealed that dual loss of NRP1 and NRP2 promotes the rapid translocation of VEGFR-2 complexes to Rab7+ late endosomes for proteosomal degradation upon acute VEGF-A_165_ stimulation, likely resulting in a severely moderated VEGFR-2 response. This work supports that of Ballmer-Hofer *et al*., who delineated that in the absence of NRP1, or in ECs stimulated with a non-NRP1-binding VEGF-A isoform, VEGFR-2 is re-routed to the degradative pathway specified by Rab7 vesicles. Importantly, this was found to occur only following 30 minutes VEGF-A stimulation [51], suggesting that the rate of VEGFR-2 degradation is accelerated when NRP1 and NRP2 are lost in tandem, as we observed changes after 5 minutes of stimulation.

In conclusion, our findings show that the activity of endothelial NRPs together is required for sustained tumour angiogenesis, and support a hypothesis that maximum anti-angiogenic efficacy can be achieved by co-targeting both NRP1 and NRP2 rather than targeting either receptor individually. Dual loss of both NRPs was found to severely abrogate tumour development and tumour angiogenesis in multiple models of cancer, in addition to secondary tumour development. This work provides strong evidence for the need to develop novel targeted therapeutics specific for both endothelial NRP1 and NRP2 receptors, against pathologies characterised by uncontrolled vascular expansion.

## Materials and Methods

### Animal generation

All experiments were performed in accordance with UK home office regulations and the European Legal Framework for the Protection of Animals used for Scientific Purposes (European Directive 86/609/EEC), prior to the start of this project. NRP1 (NRP1^flfl^) [52] and NRP2 floxed (NRP2^flfl^) [27] mice were purchased from Jackson Laboratories (Bar Harbour, Maine, USA. All animals were bred on a pure C57/BL6 background.

The PCR analysis to confirm floxing was carried out using the following oligonucleotide primers: Forward NRP1 primer: *5’ -AGGTTAGGCTTCAGGCCAAT-3’*, Reverse NRP1 primer: *5’-GGTACCCTGGGTTTTCGATT-3’*. Forward NRP2 (WT reaction) primer (Reaction A): *5’-CAGGTGACTGGGGATAGGGTA-3’*, common NRP2 primer (Reaction A + B): *5’-AGCTTTTGCCTCAGGACCCA-3’*, forward NRP2 primer (^flfl^ reaction) (Reaction B): *5’-CCTGACTACTCCCAGTCATAG -3*’.

Transgenic mice expressing a tamoxifen-inducible PDGFb-iCreER^T2^ allele in vascular ECs were provided by Marcus Fruttiger (UCL, London, UK). PCR confirmation of Cre-recombinase status was performed using the following oligonucleotide primers: Forward primer: *5’-GCCGCCGGGATCACTCTC-3’*, Reverse primer: *5’-CCAGCCGCCGTCGCAACT-3’*. NRP1^flfl^ and NRP2^flfl^ mice were bred with PDGFb.iCreER^T2^ mice to generate NRP1^flfl^.Pdgfb-iCreER^T2^ and NRP2^flfl^.PDGFb.iCreER animals. NRP1^flfl^.Pdgfb-iCreER^T2^ and NRP2^flfl^.PDGFb.iCreER mice were subsequently bred to generate NRP1^flfl^;NRP2^flfl^.Pdgfb-iCreER^T2^ animals. PDGFβ-iCreER^T2^ expression was maintained exclusively on breeding males to ensure the production of both Cre-negative and positive offspring, and therefore the use of littermate controls.

### CMT19T tumour growth assays

Mice received intraperitoneal (IP) injections of tamoxifen (75 mg/kg bodyweight, 2mg/ml stock in corn oil) thrice weekly for the duration of the experiment from D-4 to D17 to induce target deletion. CMT19T lung carcinoma cells (CR-UK Cell Production) (1×10^6^) were implanted subcutaneously (SC) into the flank of mice at D0 and allowed to develop until D18. On D18, mice were killed, and tumour volumes and weights measured. Tumour volume was calculated according to the formula: length x width^2^ x 0.52.

### Intervention tumour growth assays

CMT19T lung carcinoma cells (1×10^6^) or B16-F10 melanoma cells (4×10^5^) were implanted subcutaneously (SC) into the flank of mice at D0 and allowed to develop until D18. PyMT-BO1 cells (1×10^5^ in matrigel) were implanted orthotopically into the inguinal mammary fat pad under anaesthesia, and allowed to develop until D15. Mice received intraperitoneal (IP) injections of tamoxifen (75 mg/kg bodyweight, 2 mg/ml stock) thrice weekly for the duration of the experiment from D7 to induce target deletion. On D18/D15, mice were killed, and tumour volumes and weights measured.

### Tissue immunofluorescence analysis

Cryopreserved tumour sections were fixed in 4% PFA for 10 minutes, then washed in PBS 0.3% triton-X100, and in PBLEC (1x PBS, 1% Tween 20, 0.1 mM CaCl_2,_ 0.1 mM MgCl_2,_ 0.1 mM MnCl_2_), before being incubated in Dako protein block serum free (Agilent). Sections were then incubated overnight at 4°C in primary antibodies against NRP1 (clone AF566; R&D), NRP2 (clone Sc-13117; Santa-Cruz Biotechnology (SCB)), endomucin (clone Sc-65495; SCB), Ki-67 (Ab15580; Abcam), EDA-FN (clone F6140; Sigma), VEGFR-2 (clone 2479; Cell signalling technologies (CST)), *p*-VEGFR-2^Y1175^ (clone 2478; CST). Following primary antibody incubation, sections were washed again in PBS 0.3% triton-X100 before being incubated in the appropriate Alexa fluor secondary antibody for 2 hours at RT. Sections were blocked in Sudan black before mounting. Sections were imaged at 20 X magnification using a Zeiss AxioImager M2 microscope (AxioCam MRm camera).

Tumour vascular density was assessed by counting the number of endomucin-positive vessels per mm^2^ from 3 representative ROIs averaged per section, subsequently averaged over 3 sections/tumour. Vascular density of lung nodules was measured using a previously described ImageJ^TM^ macro application [53].

### Cell isolation, immortalisation, and cell culture

Primary mouse lung microvascular endothelial cells (mLMECs) were isolated from WT C57/BL6 adult mice. mLMECs were twice subject to magnetic activated cell sorting (MACS) to positively select for endomucin^+^ ECs as previously described by Reynolds & Hodivala-Dilke [54]. ECs were immortalised using polyoma-middle-T-antigen (PyMT) by retroviral transfection as previously described by Robinson et al [55]. Immortalised mLMECs were cultured in media composed of a 1:1 mix of Ham’s F-12:DMEM (low glucose) medium supplemented with 10% FBS, 100 units/mL penicillin/streptomycin (P/S), 2 mM glutamax, 50 μg/mL heparin (Sigma). ECs were grown at 37°C in a humidified incubator with 5% CO_2_ unless otherwise stated. For experimental analyses, plasticware was coated using 10 µg/ml human plasma fibronectin (FN) (Millipore).

EC stimulation was achieved using 30 ng/ml VEGF-A_164_ (VEGF-A) (mouse equivalent of VEGF-A_165_) post 3 hours incubation in serum-free medium (OptiMEM^®^; Invitrogen). VEGF-A was made in-house as previously described by Krilleke et al [56].

### siRNA transfection

Immortalised ECs were transfected with non-targeting control siRNA (*si*Ctrl) or mouse-specific siRNA constructs against NRP1 or NRP2 (Dharmacon), suspended in nucleofection buffer (200 mM Hepes, 137 mM NaCl, 5 mM KCl, 6 mM D-glucose, and 7 mM Na_2_HPO_4_ in nuclease-free water; filter sterilised). Nucleofection was performed using the Amaxa 4D-nucleofector system (Lonza) using program EO-100 according to manufacturer’s instructions.

### VEGF-A_165_ stimulation

ECs were incubated in serum-free OptiMEM^®^ for 3 hours prior to VEGF-A_165_ stimulation (30 ng/ml) for the indicated timepoints. ECs were then immediately placed on ice, washed twice with ice-cold PBS, then lysed in electrophoresis sample buffer (ESB) (Tris-HCL: 65 mM pH 7.4, sucrose: 60 mM, 3 % SDS).

### Deoxycholate (DOC) Buffer-extraction

Following VEGF-A_165_ stimulation, ECs were lysed in DOC lysis buffer (20 mM Tris, pH 8.5, 1% sodium deoxycholate, 2 mM iodoacetamide, 2 mM EDTA) in the presence of 100X Halt protease inhibitor cocktail, cleared by centrifugation, and the insoluble fraction isolated. Soluble and insoluble fractions were separated by SDS-PAGE and subjected to Western blot analysis.

### Western blotting

Equivalent protein concentrations were loaded onto 8% polyacrylamide gels and subjected to SDS-PAGE. Proteins were transferred to a nitrocellulose membrane (Sigma) before being incubated in 5 % milk powder. Membranes were then incubated overnight in primary antibody diluted 1:1000 at 4 °C. Membranes were washed with 0.1% Tween-20 in PBS (PBST) and incubated in an appropriate horseradish peroxidase (HRP)-conjugated secondary antibody (Dako) diluted 1:2000 for 2 hours at RT. Bands were visualised by incubation with a 1:1 solution of Pierce ECL Western Blotting Substrate (Thermo). Chemiluminescence was detected on a ChemiDoc^TM^ MP Imaging System (BioRad). Densitometric readings of band intensities were obtained using ImageJ^TM^. Primary antibodies were all purchased from CST unless otherwise stated: *p*-VEGFR-2^Y1175^ (clone 2478; CST), VEGFR-2 (2479), NRP2 (3326), NRP1 (3725), HSC70 (clone B-6; SCB), EDA-FN (F6140; Sigma).

### Metastasis experiments

luciferase^+^-tagged B16-F10 melanoma cells (1×10^6^) were intravenously (IV) injected into the tail vein of mice at D0 and allowed to disseminate until D14. Mice received intraperitoneal (IP) injections of tamoxifen (75 mg/kg bodyweight, 2mg/ml stock) thrice weekly for the duration of the experiment from D3 to induce target deletion. On D14, mice were killed, and lungs removed for bioluminescence imaging and subsequent immunofluorescence analysis of sections.

### Biotin-surface protein labelling

ECs were washed twice on ice with Soerensen buffer (SBS) pH 7.8 (14.7mM KH_2_PO_4_, 2mM Na_2_HPO_4_, and 120mM Sorbitol pH 7.8). Surface proteins were labelled with 0.3 mg/ml biotin (Thermo Scientific) in SBS for 30 minutes at 4 °C. Unreacted biotin was quenched in 100 mM glycine for 10 minutes. Biotin stripping was achieved by incubation with 100 mM MESNA (Sigma) for 75 minutes at 4 °C. Unreacted MESNA was quenched with 100 mM iodoacetamide (Sigma) for 10 minutes. ECs were lysed in lysis buffer (25 mM Tris-HCl, pH 7.4, 100 mM NaCl, 2 mM MgCl_2_, 1 mM Na_3_VO_4_, 0.5 mM EGTA, 1% Triton X-100, 5% glycerol, and protease inhibitors), and placed on ice as described previously [34]. Lysates were cleared by centrifugation at 12,000g for 20 minutes at 4°C, then quantified using the DC BioRad protein assay. Equivalent protein concentrations were immunoprecipitated with Protein G Dynabeads^TM^ (Invitrogen) coupled to a mouse anti-biotin primary antibody. Immunoprecipitated biotin-labelled proteins were separated by SDS-PAGE and subjected to Western blot analysis.

### Immunocytochemistry

ECs were seeded onto acid-washed, oven sterilised glass coverslips for 3 hours. Following VEGF-A stimulation, ECs were fixed in 4 % paraformaldehyde, washed in PBS, blocked and permeabilised with 10% goat serum in PBS 0.3% triton X-100. ECs were incubated in primary antibody diluted 1:100 overnight at 4°C. Coverslips were then PBS washed and incubated in an appropriate Alexa fluor secondary antibody diluted 1:200 in PBS for 1 hour at RT. Coverslips were mounted using flouromount G with DAPI^TM^. Images were captured using a Zeiss AxioImager M2 microscope (AxioCam MRm camera) at 63x magnification. Primary antibodies: VEGFR-2 (CST; 2479), Rab7 (CST; 17286).

### Statistical analysis

The graphic illustrations and analyses to determine statistical significance were generated using GraphPad Prism 9 software and Student’s t-tests unless otherwise stated. Statistical analysis between Cre-positive groups was performed using one-way ANOVA tests. Bar charts show mean values and the standard error of the mean (+SEM). Asterisks indicate the statistical significance of p values: p > 0.05 = NS (not significant), *p < 0.05, **p < 0.01, ***p < 0.001 and ****p < 0.0001.

## Acknowledgments

This work was supported by funding from: the UKRI Biotechnology and Biological Sciences Research Council (BBSRC) Norwich Research Park (NRP) Biosciences Doctoral Training Partnership (DTP) (grant numbers BB/M011216/1, BB/J014524/1), and a BigC studentship to SDR/CAP (18-15R). Robinson is also partially funded by the BBSRC Institute Strategic Programme Gut Microbes and Health BB/R012490/1 and its constituent project (BBS/E/F/000PR10355). Additionally, we thank Norfolk Fundraisers, Mrs Margaret Doggett, and the Colin Wright Fund for their kind support and fundraising over the years.

## Data availability statement

The raw data supporting the conclusions of this manuscript will be made available by the authors, without undue reservation, to any qualified researcher.

## Competing interests

All authors declare no conflicts of interest.

**Supplementary Figure 1:**
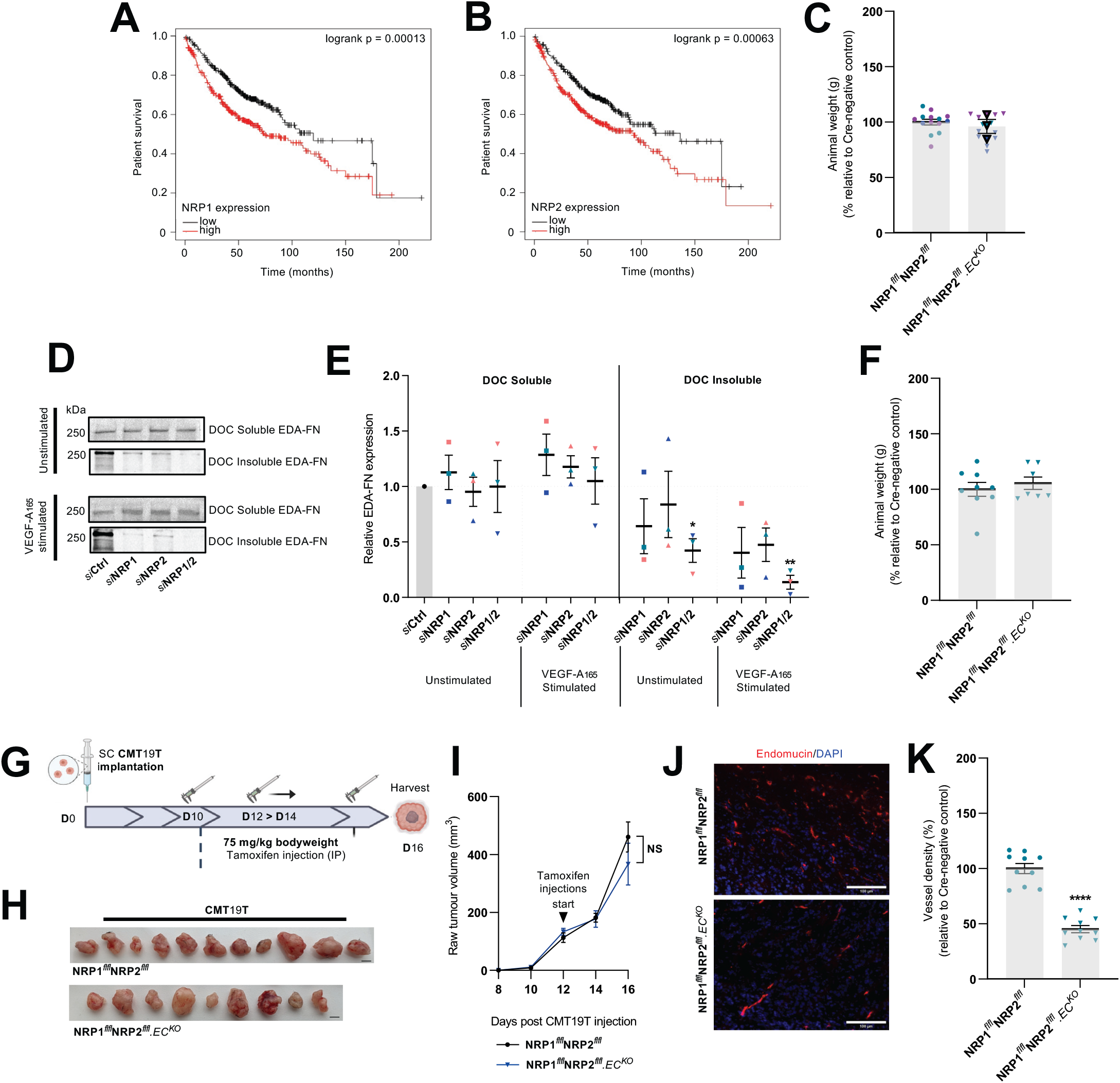
A) Determination of prognostic value of NRP1 and NRP2 receptor mRNA expression in lung carcinoma patients (n = 719) using www.kmplot.com. Kaplan-Meier survival plot of lung carcinoma patients with high NRP1 mRNA expression (*Affymetrix ID: 210615_at*). **B**) Kaplan-Meier survival plot of lung carcinoma patients with high NRP2 mRNA expression (*Affymetrix ID: 214632_at*). Respective logrank *p* values are shown. **C**) Quantification of mean animal weight measured at point of harvest. Data presented as a percentage of the average animal weight observed in respective littermate controls. Error bars show mean ± SEM, N=3 (n≥12). **D**) siRNA-treated ECs were seeded onto 10 µg/ml FN for 48 hours, then incubated in serum-free OptiMEM for 3 hours. ECs were subject to 5 minutes of stimulation with 30 ng/ml VEGF-A before being washed twice on ice with PBS and lysed. Lysates were cleared by centrifugation for 30 minutes at 4 °C, allowing for the isolation of soluble and insoluble fractions. Soluble and insoluble fractions were quantified using the DC protein assay, separated by SDS-PAGE and subjected to Western blot analysis. Membranes were incubated in anti-EDA-FN primary antibody. Panels show representative levels of EDA-FN expression quantified from soluble and insoluble fractions, in the presence or absence of VEGF-A stimulation. **E**) Quantification of relative EDA-FN expression. Quantification shows mean densitometric analysis obtained using ImageJ^TM^. Error bars show mean ± SEM; N=3. **F**) Quantification of mean animal weight measured at point of harvest. Data presented as a percentage of the average animal weight observed in respective littermate controls. Error bars show mean ± SEM, n≥6. **G**) Delayed experimental schematic: tamoxifen-induced activation of Cre-recombinase and thus deletion of targets was employed via the following regime. Cre-positive and Cre-negative littermate control mice received intraperitoneal (IP) injections of tamoxifen (75 mg/kg bodyweight, 2mg/ml stock) on D12 and D14 to induce Cre-recombinase activity. CMT19T lung carcinoma cells (1×10^6^) were implanted subcutaneously (SC) into the flank of mice at D0 and allowed to grow until D16. **H**) Representative images of CMT19T tumours harvested on D16 removed from Cre-negative and positive mice. Scale bar shows 5 mm. **I**) Raw tumour volume growth kinetics from 10 days post CMT19T injections to harvest. Error bars show mean ± SEM; n≥9. **J**) Representative tumour sections from Cre-negative and Cre-positive tumours showing endomucin^+^ blood vessels. Scale bar = 100 µm. **K**) Quantification of % blood vessel density per mm^2^ from CMT19T tumours. Mean quantification performed on 3x ROIs per tumour section, from 1-3 sections per tumour. Data presented as a percentage of the average % vessel density observed in their Cre-negative littermate controls. Error bars show mean ± SEM; n≥9. Asterixis indicate significance.

**Supplementary Figure 2:**
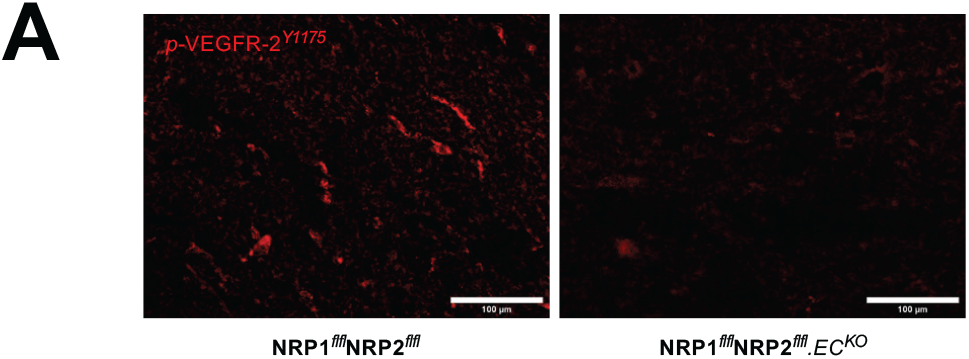
A) Representative tumour sections from Cre-negative and Cre-positive tumours showing *p*-VEGFR-2Y^1175^ expression. Scale bar = 100 µm.

## Notes

### Competing Interest Statement

The authors have declared no competing interest.

## References

1. J. Folkman, “What is the evidence that tumors are angiogenesis dependent?,” J. Natl. Cancer Inst., vol. 82, no. 1, pp. 4–7, 1990, doi: 10.1093/jnci/82.1.4.

2. D. Hanahan and R. A. Weinberg, “Hallmarks of cancer: The next generation,” Cell, vol. 144, no. 5, pp. 646–674, 2011, doi: 10.1016/j.cell.2011.02.013.

3. A. Dumond and G. Pagès, “Neuropilins, as Relevant Oncology Target: Their Role in the Tumoral Microenvironment,” Front. Cell Dev. Biol., vol. 8, no. July, pp. 1–10, 2020, doi: 10.3389/fcell.2020.00662.

4. A. G. Sorensen et al., “Increased survival of glioblastoma patients who respond to antiangiogenic therapy with elevated blood perfusion,” Cancer Res., vol. 72, no. 2, pp. 402–407, 2012, doi: 10.1158/0008-5472.CAN-11-2464.

5. R. T. Tong, Y. Boucher, S. V. Kozin, F. Winkler, D. J. Hicklin, and R. K. Jain, “Vascular normalization by vascular endothelial growth factor receptor 2 blockade induces a pressure gradient across the vasculature and improves drug penetration in tumors,” Cancer Res., vol. 64, no. 11, pp. 3731–3736, 2004, doi: 10.1158/0008-5472.CAN-04-0074.

6. H. G. Augustin and G. Y. Koh, “Antiangiogenesis: Vessel Regression, Vessel Normalization, or Both?,” Cancer Res., vol. 82, no. 1, pp. 15–17, 2022, doi: 10.1158/0008-5472.can-21-3515.

7. T. R. Wilson et al., “Widespread potential for growth-factor-driven resistance to anticancer kinase inhibitors,” Nature, vol. 487, no. 7408, pp. 505–509, 2012, doi: 10.1038/nature11249.Widespread.

8. S. Soker, S. Takashima, H. Q. Miao, G. Neufeld, and M. Klagsbrun, “Neuropilin-1 is expressed by endothelial and tumor cells as an isoform-specific receptor for vascular endothelial growth factor,” Cell, vol. 92, no. 6, pp. 735–745, 1998, doi: 10.1016/S0092-8674(00)81402-6.

9. I. Zachary, “Neuropilins: Role in Signalling, Angiogenesis and Disease,” Chem Immunol Allergy, vol. 99, pp. 37–70, 2014.

10. L. Zhao, H. Chen, L. Lu, L. Wang, X. Zhang, and X. Guo, “New insights into the role of co-receptor neuropilins in tumour angiogenesis and lymphangiogenesis and targeted therapy strategies,” J. Drug Target., vol. 29, no. 2, pp. 155–167, 2021, doi: https://doi.org/10.1080/1061186X.2020.1815210.

11. A. Dumond and G. Pagès, “Relevance of Neuropilin 1 and Neuropilin 2 Targeting for Cancer Treatment,” J. Cancer Immunol., vol. 3, no. 2, pp. 111–114, 2021, doi: 10.33696/cancerimmunol.3.046.

12. B. Favier et al., “Neuropilin-2 interacts with VEGFR-2 and VEGFR-3 and promotes human endothelial cell survival and migration,” Blood, vol. 108, no. 4, pp. 1243–1250, 2006, doi: 10.1182/blood-2005-11-4447.

13. T. Kawakami et al., “Neuropilin 1 and neuropilin 2 co-expression is significantly correlated with increased vascularity and poor prognosis in nonsmall cell lung carcinoma,” Cancer, vol. 95, no. 10, pp. 2196–2201, 2002, doi: 10.1002/cncr.10936.

14. L. M. Ellis, “The role of neuropilins in cancer,” Mol. Cancer Ther., vol. 5, no. 5, pp. 1099–1107, 2006, doi: 10.1158/1535-7163.MCT-05-0538.

15. D. R. Bielenberg, Y. Hida, A. Shimizu, A. Kaipainen, C. C. Kreuter MKim, and M. Klagsbrun, “Semaphorin 3F, a chemorepulsant for endothelial cells, induces a …,” J Clin Invest, vol. 114, no. 9, pp. 1260–1271, 2004, doi: 10.1172/JCI200421378.1260.

16. A. Vales et al., “Myeloid leukemias express a broad spectrum of VEGF receptors including neuropilin-1 (NRP-1) and NRP-2,” Leuk. Lymphoma, vol. 48, no. 10, pp. 1997–2007, 2007, doi: 10.1080/10428190701534424.

17. R. E. Bachelder, G. Robinson, J. Chung, M. A. Wendt, L. M. Shaw, and A. M. Mercurio, “Vascular endothelial growth factor is an autocrine survival factor for neuropilin-expressing breast carcinoma cells,” Cancer Res., vol. 61, no. 15, pp. 5736–5740, 2001.

18. S. Lantuéjoul, B. Constantin, H. Drabkin, C. Brambilla, J. Roche, and E. Brambilla, “Expression of VEGF, semaphorin SEMA3F, and their common receptors neuropilins NP1 and NP2 in preinvasive bronchial lesions, lung tumours, and cell lines,” J. Pathol., vol. 200, no. 3, pp. 336–347, 2003, doi: 10.1002/path.1367.

19. Y. Cao et al., “Neuropilin-2 promotes extravasation and metastasis by interacting with endothelial α5 integrin,” Cancer Res., vol. 73, no. 14, pp. 4579–4590, 2013, doi: 10.1158/0008-5472.CAN-13-0529.

20. U. Yaqoob et al., “Neuropilin-1 stimulates tumor growth by increasing fibronectin fibril assembly in the tumor microenvironment,” Cancer Res., vol. 72, no. 16, pp. 4047–4059, 2012, doi: 10.1158/0008-5472.CAN-11-3907.

21. C. Binetruy-Tournaire, Roselyne Demangel et al., “Identification of a peptide blocking vascular endothelial growth factor (VEGF)-mediated angiogenesis,” EMBO J., vol. 19, no. 7, pp. 1525–1533, 2000.

22. A. Jarvis et al., “Small molecule inhibitors of the neuropilin-1 vascular endothelial growth factor A (VEGF-A) interaction,” J. Med. Chem., vol. 53, no. 5, pp. 2215–2226, 2010, doi: 10.1021/jm901755g.

23. M. Srichai and R. Zent, Integrin Structure and Function: Cell-Extracellular Matrix Interactions in Cancer. Springer, Boston, MA, 2010.

24. L. Borriello et al., “Structure-based discovery of a small non-peptidic Neuropilins antagonist exerting in vitro and in vivo anti-tumor activity on breast cancer model,” Cancer Lett., vol. 349, no. 2, pp. 120–127, 2014, doi: 10.1016/j.canlet.2014.04.004.

25. W. Q. Liu et al., “NRPa-308, a new neuropilin-1 antagonist, exerts in vitro antiangiogenic and anti-proliferative effects and in vivo anti-cancer effects in a mouse xenograft model,” Cancer Lett., vol. 414, pp. 88–98, 2018, doi: 10.1016/j.canlet.2017.10.039.

26. A. Dumond et al., “Neuropilin 1 and Neuropilin 2 gene invalidation or pharmacological inhibition reveals their relevance for the treatment of metastatic renal cell carcinoma,” J. Exp. Clin. Cancer Res., vol. 40, no. 1, pp. 1–18, 2021, doi: 10.1186/s13046-021-01832-x.

27. A. Walz, I. Rodriguez, and P. Mombaerts, “Aberrant Sensory Innervation of the Olfactory Bulb in Neuropilin-2 Mutant Mice,” J. Neurosci., vol. 22, no. 10, pp. 4025–4035, 2002, doi: 10.1523/jneurosci.22-10-04025.2002.

28. S. Claxton, V. Kostourou, S. Jadeja, P. Chambon, K. Hodivala-Dilke, and M. Fruttiger, “Efficient, inducible cre-recombinase activation in vascular endothelium,” Genesis, vol. 46, no. 2, pp. 74–80, 2008, doi: 10.1002/dvg.20367.

29. C. J. Benwell, J. A. G. E. Taylor, and S. D. Robinson, “Endothelial neuropilin-2 influences angiogenesis by regulating actin pattern development and α5-integrin-p-FAK complex recruitment to assembling adhesion sites,” FASEB J., vol. 35, no. 8, pp. 1–19, 2021, doi: 10.1096/fj.202100286R.

30. A. Lanczky and B. Gyorffy, “Web-based survival analysis tool tailored for medical research (KMplot): development and implementation,” J Med Internet Res, vol. 23, no. 7, 2021, doi: 10.2196/27633.

31. S. Gopal et al., “Fibronectin-guided migration of carcinoma collectives,” Nat. Commun., vol. 8, 2017, doi: 10.1038/ncomms14105.

32. R. N. Kaplan et al., “VEGFR1-positive haematopoietic bone marrow progenitors initiate the pre-metastatic niche,” Nature, vol. 438, no. 7069, pp. 820–827, 2005, doi: 10.1038/nature04186.VEGFR1-positive.

33. H. Kumra and D. P. Reinhardt, “Fibronectin-targeted drug delivery in cancer,” Adv. Drug Deliv. Rev., vol. 97, no. April, pp. 101–110, 2016, doi: 10.1016/j.addr.2015.11.014.

34. D. Valdembri et al., “Neuropilin-1/GIPC1 signaling regulates α5β1 integrin traffic and function in endothelial cells,” PLoS Biol., vol. 7, no. 1, 2009, doi: 10.1371/journal.pbio.1000025.

35. X. Su et al., “Antagonizing integrin beta3 increases immune suppression in cancer,” vol. 76, no. 12, pp. 3484–3495, 2017, doi: 10.1158/0008-5472.CAN-15-2663.Antagonizing.

36. M. Potez et al., “Characterization of a B16-F10 melanoma model locally implanted into the ear pinnae of,” pp. 1–19, 2018.

37. L. Zhang, J. J. Hu, and F. Gong, “MG132 inhibition of proteasome blocks apoptosis induced by severe DNA damage,” Cell Cycle, vol. 10, no. 20, pp. 3515–3518, 2011, doi: 10.4161/cc.10.20.17789.

38. L. Zhang et al., “MG132-mediated inhibition of the ubiquitin-proteasome pathway ameliorates cancer cachexia,” J. Cancer Res. Clin. Oncol., vol. 139, no. 7, pp. 1105–1115, 2013, doi: 10.1007/s00432-013-1412-6.

39. F. Jin, D. Xiao, T. Zhao, and M. Yu, “Proteasome inhibitor MG132 suppresses pancreatic ductal adenocarcinoma-cell migration by increasing ESE3 expression,” Oncol. Lett., vol. 19, no. 1, pp. 858–868, 2020, doi: 10.3892/ol.2019.11157.

40. M. Hein and S. Graver, “Tumor cell response to bevacizumab single agent therapy in vitro,” Cancer Cell Int., vol. 13, no. 1, pp. 1–11, 2013, doi: 10.1186/1475-2867-13-94.

41. S. D. Liu, L. P. Zhong, J. He, and Y. X. Zhao, “Targeting neuropilin-1 interactions is a promising anti-Tumor strategy,” Chin. Med. J. (Engl)., vol. 134, no. 5, pp. 508–517, 2021, doi: 10.1097/CM9.0000000000001200.

42. K. Pal, V. S. Madamsetty, S. K. Dutta, E. Wang, R. S. Angom, and D. Mukhopadhyay, “Synchronous inhibition of mTOR and VEGF/NRP1 axis impedes tumor growth and metastasis in renal cancer,” npj Precis. Oncol., vol. 3, no. 1, pp. 1–11, 2019, doi: 10.1038/s41698-019-0105-2.

43. J. Powell et al., “Small Molecule Neuropilin - 1 Antagonists Combine Antiangiogenic and Antitumor Activity with Immune Modulation through Reduction of Transforming Growth Factor Beta (TGF β) Production in Regulatory T - Cells,” 2018, doi: 10.1021/acs.jmedchem.8b00210.

44. Z. Huang et al., “NRP1 promotes cell migration and invasion and serves as a therapeutic target in nasopharyngeal carcinoma,” Int. J. Clin. Exp. Pathol., vol. 11, no. 5, p. 2460, 2018.

45. L. Wang et al., “N2E4, a Monoclonal Antibody Targeting Neuropilin-2, Inhibits Tumor Growth and Metastasis in Pancreatic Ductal Adenocarcinoma via Suppressing FAK/Erk/HIF-1α Signaling,” Front. Oncol., vol. 11, no. July, pp. 1–12, 2021, doi: 10.3389/fonc.2021.657008.

46. M. Caunt et al., “Blocking Neuropilin-2 Function Inhibits Tumor Cell Metastasis,” Cancer Cell, vol. 13, no. 4, pp. 331–342, 2008, doi: 10.1016/j.ccr.2008.01.029.

47. X. Li et al., “Nordihydroguaiaretic acid impairs prostate cancer cell migration and tumor metastasis by suppressing neuropilin 1,” Oncotarget, vol. 7, no. 52, pp. 86225–86238, 2016, doi: 10.18632/oncotarget.13368.

48. G. Efthymiou, A. Saint, M. Ruff, Z. Rekad, D. Ciais, and E. Van Obberghen-Schilling, “Shaping Up the Tumor Microenvironment With Cellular Fibronectin,” Front. Oncol., vol. 10, no. April, pp. 1–18, 2020, doi: 10.3389/fonc.2020.00641.

49. X. Su et al., “FN-EDA mediates angiogenesis of hepatic fibrosis via integrin-VEGFR2 in a CD63 synergetic manner,” Cell Death Discov., vol. 6, no. 1, 2020, doi: 10.1038/s41420-020-00378-9.

50. L. Xiang, G. Xie, J. Ou, X. Wei, F. Pan, and H. Liang, “The extra domain a of fibronectin increases VEGF-C expression in colorectal carcinoma involving the PI3K/AKT signaling pathway,” PLoS One, vol. 7, no. 4, pp. 1–10, 2012, doi: 10.1371/journal.pone.0035378.

51. K. Ballmer-Hofer, A. E. Andersson, L. E. Ratcliffe, and P. Berger, “Neuropilin-1 promotes VEGFR-2 trafficking through Rab11 vesicles thereby specifying signal output,” Blood, vol. 118, no. 3, pp. 816–826, 2011, doi: 10.1182/blood-2011-01-328773.

52. C. Gu et al., “Neuropilin-1 conveys semaphorin and VEGF signaling during neural and cardiovascular development,” Dev. Cell, vol. 5, no. 1, pp. 45–57, 2003, doi: 10.1016/S1534-5807(03)00169-2.

53. J. Lambert et al., “ADAMTS-1 and syndecan-4 intersect in the regulation of cell migration and angiogenesis,” J. Cell Sci., vol. 133, no. 7, pp. 1–15, 2020, doi: 10.1242/jcs.235762.

54. L. Reynolds and K. Hodivala-Dilke, “Primary mouse endothelial cell culture for assays of angiogenesis.,” Methods Mol Med, vol. 120, pp. 503–509, 2006.

55. S. D. Robinson et al., “?vβ3 Integrin Limits the Contribution of Neuropilin-1 To Vascular Endothelial Growth Factor-Induced Angiogenesis,” J. Biol. Chem., vol. 284, no. 49, pp. 33966–33981, 2009, doi: 10.1074/jbc.M109.030700.

56. D. Krilleke et al., “Molecular mapping and functional characterization of the VEGF164 heparin-binding domain,” J. Biol. Chem., vol. 282, no. 38, pp. 28045–28056, 2007, doi: 10.1074/jbc.M700319200.

